# Resting Galvanic Skin Response Reflects Fluctuations in Creativity Potential for Solving Creativity Tasks

**DOI:** 10.64898/2026.08.23.746567

**Authors:** Tzu-Ling Liu, Miyoko Street, Zenas C. Chao

**Affiliations:** International Research Center for Neurointelligence (WPI-IRCN), UTIAS, The University of Tokyo, Japan

**Keywords:** Creativity, AUT, FIT, biophysiological measurements, EDA (GSR)

## Abstract

Creative performance fluctuates from moment to moment, suggesting that it depends partly on transient internal states present before creative thinking begins. Although such fluctuations have been identified in central neural activity, it remains unclear whether they are also reflected in autonomic physiology. We examined whether cardiac and electrodermal activity during a brief pre-trial resting period was associated with subsequent creative performance. Photoplethysmography, electrocardiography, and electrodermal activity were recorded while 28 participants completed the Alternative Uses Test and Fusion Innovation Test, assessing divergent and convergent creative thinking, respectively. Data from 27 participants was included in the analyses. Heart rate, heart rate variability, tonic skin conductance level, and phasic skin conductance activity were extracted from observation windows ranging from 11 to 30 s within a 30-s pre-trial rest period. Trial-level creativity was evaluated using GPT-based ratings of novelty, feasibility, and goal attainment. Linear mixed-effects models showed that higher pre-trial tonic skin conductance level was consistently associated with better subsequent creative performance across tasks, with overall model fit peaking at a 21-s observation window. Permutation-based feature-importance analysis provided convergent support for the contribution of tonic skin conductance, whereas the cardiac and phasic electrodermal indicators showed no reliable independent associations. However, the model explained only a small proportion of behavioral variance. These findings suggest that tonic sympathetic arousal reflects a momentary physiological state associated with creativity potential, while autonomic signals alone remain insufficient for accurate individual-level prediction.

**AI Use:** In the current study, generative artificial intelligence (AI) tools were utilized to assist with code development during data analysis, reference organization, and proofreading during manuscript preparation. The core experimental design, analytical methodologies, and primary drafting of the manuscript were performed entirely by the authors. All AI-generated code and text edits were thoroughly reviewed and verified by the authors to ensure that the analysis strictly adhered to the designated pipeline, parameter selections were theoretically sound and properly referenced, and the final text accurately reflected the original conceptual intent.

## Introduction

Creative performance is often treated as a relatively stable individual ability, but it also fluctuates from moment to moment within the same person. These fluctuations suggest that performance depends partly on transient internal states present before a creative task begins. Using EEG data from the same experiment, we previously identified a resting-state functional connectivity pattern, termed the Creativity Potential Network, that was associated with subsequent performance across divergent- and convergent-thinking tasks (Chao, Street, and Liu 2025). This finding raises a further question: whether momentary fluctuations in creativity potential are also reflected in peripheral physiological activity.

Peripheral physiological signals are regulated by the autonomic nervous system (ANS) and provide accessible measures of bodily arousal. The Dual Pathway to Creativity Model proposes that creative performance can arise through cognitive flexibility or persistence, both of which are influenced by activation level and affective state (De Dreu, Baas, and Nijstad 2008). Empirical studies have also linked autonomic activity to creative cognition. In a compound remote associates task, electrodermal activity immediately before solution reporting was higher for problems solved through insight than for those solved analytically (Shen et al. 2018). In a divergent-thinking task, lower low-frequency heart rate variability was associated with greater response fluency (Loudon and Deininger 2016). Together, these findings indicate that autonomic activity is related to creative performance.

However, existing evidence does not establish whether autonomic activity reflects a pre-existing state of creative readiness or merely accompanies the creative process itself. Previous studies have primarily examined physiological activity during task engagement or immediately before a response, when it may already be shaped by task demands, cognitive effort, emerging solution recognition, or response preparation. The critical unresolved question is therefore whether spontaneous, task-free autonomic activity before task onset is associated with subsequent creative performance. Demonstrating such a relationship would extend the concept of state-dependent creativity from central neural activity to peripheral physiology. It would also have practical value, because cardiac and electrodermal signals can be recorded continuously using wearable devices, providing a more accessible approach than fMRI or EEG for monitoring momentary creativity-related states.

To address this question, we tested whether autonomic activity during a 30-s pre-trial resting period was associated with subsequent creative performance. Twenty-eight participants completed the Alternative Uses Test (AUT, Guilford et al. 1960), assessing divergent thinking, and the Fusion Innovation Test (FIT, Wu et al. 2025), assessing convergent, goal-directed creative problem solving. We examined cardiac and electrodermal indicators, including heart rate, heart rate variability, tonic skin conductance level, and phasic skin conductance activity, across observation windows ranging from 11 to 30 s. Responses were evaluated for novelty, feasibility, and goal attainment using GPT-based procedures previously shown to align with human ratings (Kern, Wu, and Chao 2024; Wu et al. 2025). We asked whether any pre-trial autonomic indicator showed a reliable association with subsequent creative performance across both tasks after accounting for individual and task-related differences. Evidence for such an association would suggest that peripheral physiology reflects a momentary state of creative readiness before creative thinking begins.

## Methods

The data analyzed in the current study were collected as part of a larger experimental protocol. While the EEG data of this experiment is already published elsewhere (Chao, Street, and Liu 2025), the current study focused on the physiological signals that have not yet been studied.

### Experiment procedures

The experiment design followed the protocol described in (Chao, Street, and Liu 2025), which the protocols were approved by the Research Ethics Committee of the University of Tokyo (No 23-27). A total of 28 native Japanese-speaking university students (17 males, 11 females, aged 18-29 years, with mean ± SD = 22 ± 2.5 years) participated in the experiment. The experiment includes two creativity tasks: the Alternative Uses Test (AUT) and the Fusion Innovation Test (FIT).

As illustrated in Figure 1A, AUT showed participants a daily object and required them to think of an alternative use beyond its original use. FIT assigned two objects and one goal; it required participants to combine the two objects to achieve the goal. We adopted a two-day experiment protocol to avoid fatigue during long tasks. In each experiment day, participants completed 15 trials of AUT and 15 trials of FIT, with a 5-min resting recording before each experiment session, and a 5-min between task break that the EEG cap was removed and participant can move their body to relax (Figure 1B). Each trial started with a 30-s pre-trial resting period. Participants were required to sit still, relax, and look at the fixation cross on the screen during the resting period. After the onset of the question, they were asked to give one solution as soon as possible. Participants were asked to press the response key within the time limits to answer the question, otherwise the question was skipped and proceeded automatically.

**Figure 1.**
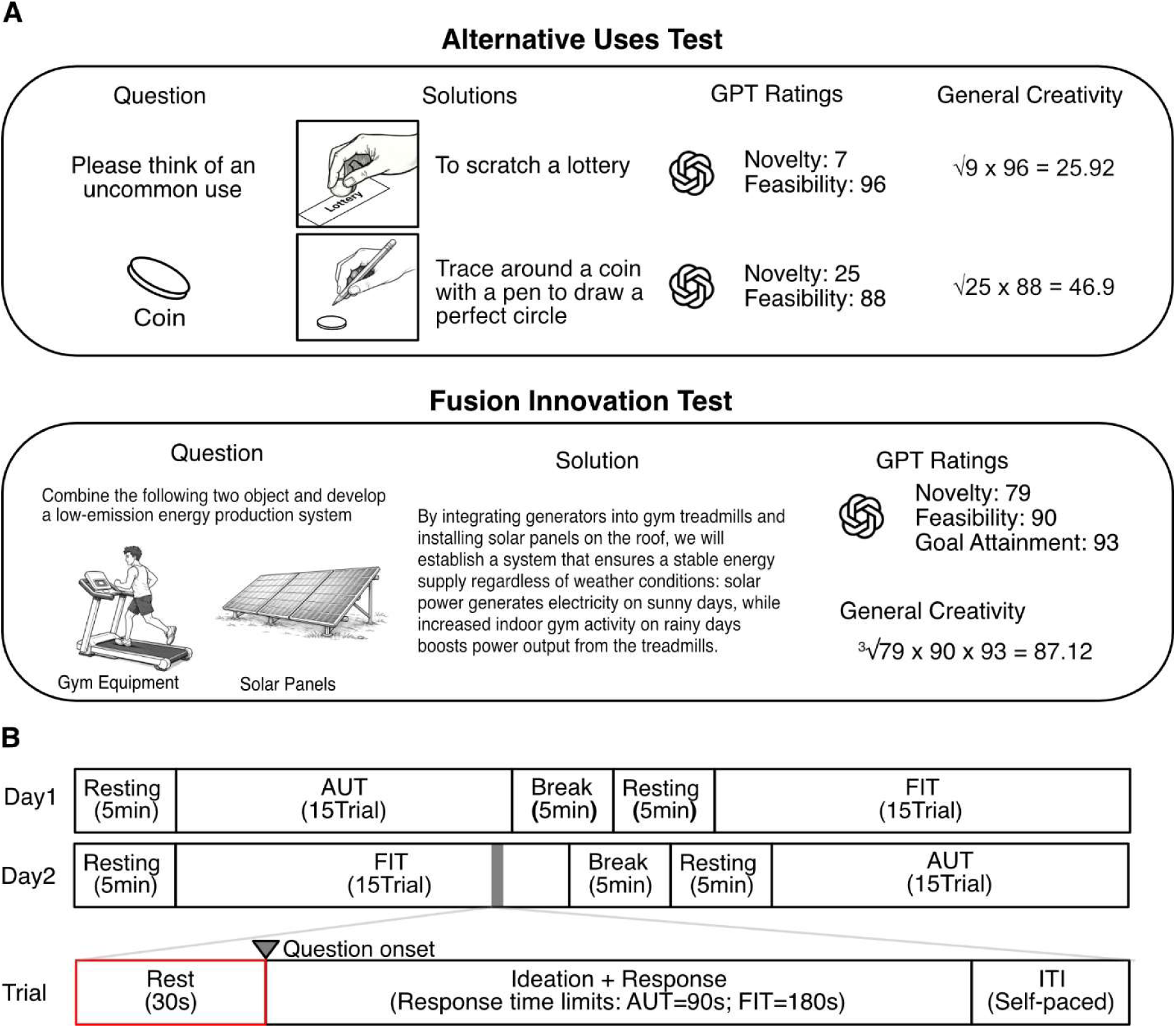
Experimental tasks and procedures. (**A**) Creativity assessment paradigms. (Top) Alternative Uses Test (AUT): Participants generate unconventional uses for everyday objects (e.g., a coin). (Bottom) Fusion Innovation Test (FIT): A novel task requiring the integration of two disparate objects to meet a specific functional goal (e.g., creating a low-emission energy system using gym equipment and solar panels). Responses are automatically scored by GPT for Novelty, Feasibility (in both AUT and FIT), and Goal Attainment (only in FIT). A General Creativity score is derived using the geometric mean of these components and subsequently z-scored within-subject. (**B**) Experimental protocol and trial structure. The study employs a 2-day counterbalanced design to control order effects. Each trial begins with a 30-s rest (red box), followed by the question onset. Response windows are 90 s for the AUT and 180 s for the FIT, followed by an inter-trial interval (ITI).

### Creativity score evaluation

Participants’ responses were merged and rated with the Large Language Model (LLM) GPT-4o, with prompts designed from previous studies (AUT: Kern, Wu, and Chao 2024; FIT: Wu et al. 2025. Based on the task and prompt design, GPT rated each AUT solution with novelty (N) and feasibility (F), and rate FIT solutions with novelty (N), feasibility (F), and goal attainment (G). All scores ranged between 1 and 100, with the higher score indicates a more novel, feasible, and better goal-attaining answer. The general creativity scores were represented by the geometric means of novelty and feasibility in AUT (*C*_AUT_ = 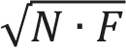), and by the geometric means of novelty, feasibility, and goal attainment in FIT (*C*_FIT_ = 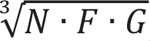). Given the distributional differences between the AUT and FIT scores, including the right-skewed distribution of FIT scores observed in previous analyses of this dataset (Chao, Street, and Liu 2025), and our focus on within-participant fluctuations in creativity, creativity scores were z-score normalized separately within each participant and task. We refer to the resulting measure as *Regularized C*:

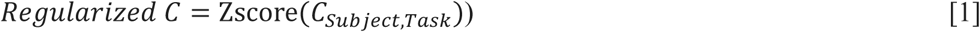

This normalization reduced variance attributable to differences between participants and tasks.

### Physiological measurement setting

The physiological data was recorded in an environment-controlled laboratory, along with EEG recording. To ensure participants’ physiological responses do not vary too much between experiment days due to biological rhythms or life routines, the two experiment days were arranged to have one-week interval. The median interval between the two sessions was 7 days, except for one participant who could not attend the second session within one week. The room temperature was maintained between 24-26 ℃ and the humidity was maintained between 40-60%. Participants were asked to follow the clothing instructions of wearing long trousers, socks to cover ankles, and short sleeve top to keep warm in the laboratory.

The current experiment collected physiological signals from the cardiovascular system, respiration system, and electrodermal activity with the wearable AIM Physiological Monitor system (Brain Products GmbH, Gilching, Germany) at a sampling rate of 500 Hz. We employed a photoplethysmogram (PPG) sensor, a pair of integrated sensors combining electrocardiogram (ECG) sensors and wireless respiration strain sensors, and a pair of electrodermal activity (GSR) sensors (see Figure 2A for sensor photos and illustration of sensor locations). The PPG sensor was clipped to the right earlobe to monitor heart rate (HR) and peripheral oxygen saturation (SpO_2_). The integrated ECG and respiration sensors were attached under the collar bones to capture cardiac activity and chest expansion. Finally, the GSR sensors were attached to the thenar and hypothenar eminences of the right palm. The GSR sensor placement was selected based on a pre-experiment test, which showed the best signal sensitivity while minimizing movement artifacts during task-related typing. Participants’ physiological responses were recorded throughout the experiment. The current study extracted the physiological signals within the 30-s resting period (red box in the last row of Figure 1B).

**Figure 2.**
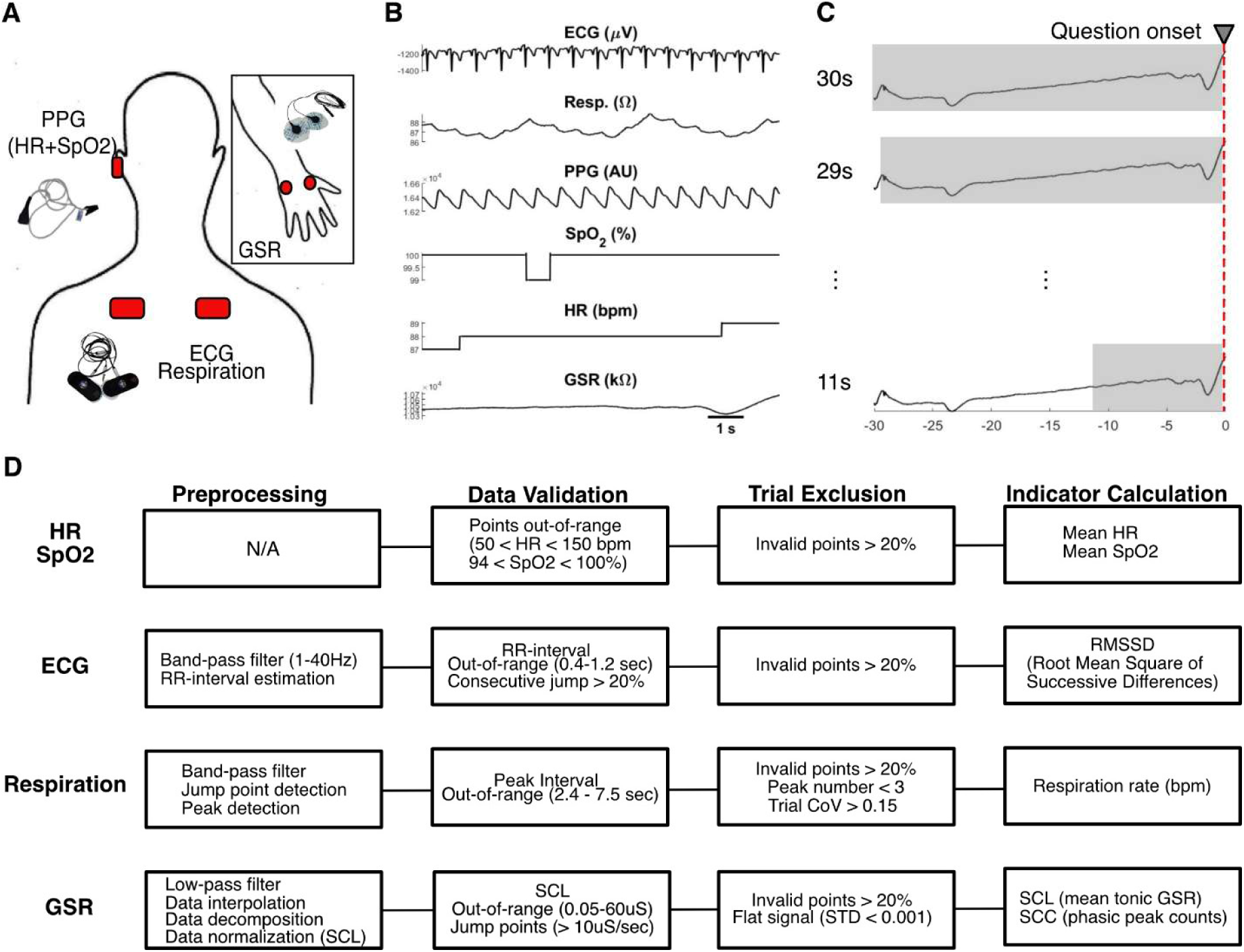
Physiological signal recording and data processing pipeline. (**A**) Sensor placement and instrumentation. Physiological signals were recorded using the CGX AIM Physiological Monitor system. A photoplethysmogram (PPG) sensor was clipped to the right earlobe to monitor heart rate (HR) and peripheral oxygen saturation (SpO2). Integrated electrocardiogram (ECG) and wireless respiration strain sensors were positioned under the collar bones to capture cardiac activity and chest expansion. Electrodermal activity (GSR) sensors were localized to the thenar and hypothenar eminences of the right palm; this placement was selected to maximize signal sensitivity while minimizing movement artifacts during task-related typing. (**B**) Representative raw traces. An illustrative 10-s data segment (Subject 001, AUT Trial 1) from all physiological channels. The vertical red dashed line indicates trial onset (T=0). (**C**) Physiological data segmentation. To identify the optimal observation window for creative performance, physiological data from the 30-s rest period are segmented into varying epoch lengths, ranging from 11 to 30 s prior to trial onset (Time 0; red dashed line). Shaded areas represent the specific data segments used for each window length analysis. (**D**) Data preprocessing pipeline consists of four primary stages: (1) Preprocessing: Signal-specific filtering and feature estimations. (2) Data validation: Identification of invalid points falling outside physical plausible ranges. (3) Trial Exclusion: Trials were rejected based on the percentage of invalid data or signal instability. For respiration and GSR, sparse artifacts in non-excluded trials were linearly interpolated to maintain signal continuity. (4) Indicator Calculation: Computation of final indicators for final statistical analysis, including Heart Rate Variability (Root Mean Square of Successive Difference, RMSSD), Respiration Rate, and GSR components (SCL represents tonic skin conductance level and SCC represent phasic skin conductance component with peak counts). For PPG-derived HR and SpO2, raw device outputs were utilized for validation and final calculation without additional filtering. For GSR signals, preprocessing and validation were mixed in order, in which definition of invalid, data interpolation and decomposition were conducted on non-normalized data, and signal normalization was only conducted on SCL signals.

### General data preprocessing

The raw data (see Figure 2B for example) from two days were merged, down-sampled to 250 Hz, and segmented into 30-s epochs immediately before the question onset. These segmented data were saved for later preprocessing. To explore the optimal length of the observation window, we dissected the 30-s data into data that covers the 11- to 30-s range before question onset (Figure 2C) and performed the processing pipeline for each data length separately. Data of each length went through a standardized analysis pipeline (Figure 2D), including preprocessing, validation, trial exclusion, and indicator calculation. During preprocessing, essential information was extracted from the raw signal. In the validation stage, data points performed out of physiological plausible range, or represent artifacts were identified and marked as invalid. In trial exclusion, trials with over 20% of invalid samples or containing certain artifact features (e.g., flat signal) were excluded from the following statistical analysis. While no universally accepted trial-level rejection criteria are available for multimodal physiological recordings, particularly for recordings as short as the 30-s intervals used here, the thresholds were defined according to the characteristics of each signal and its derived measure. Indicators were then calculated from the remaining trials and then transferred to within-subject, within-task z-scores, except for the GSR signals which were within-trial normalized before index calculation.

### Heart rate and peripheral oxygen saturation

HR and SpO_2_ signals were directly extracted from readings of the PPG sensor. Data points that fell out of the physiologically plausible range (HR: 50-150 bpm, SpO_2_: 94-100%) were marked as invalid. Trials in which more than 30% of samples were invalid were excluded. The mean values were calculated for each surviving trial.

### Heart rate variability

The ECG signals were used for estimating heart rate variability (HRV), which is an index of cardiac sympathetic and parasympathetic neural system. The raw signal was filtered between 1-40 Hz. The RR-intervals were calculated with the MATLAB function *pan_tompkins.m,* (Sedghamiz 2014; Pan and Tompkins 1985). RR intervals, time intervals between two successive R peaks in the ECG signal, were identified as abnormal if outside the plausible physiological range of 0.4 to 1.2 s (50–150 bpm) or differed from the preceding beat by more than 20% (Karey et al. 2019). If more than 20% of the total intervals within the time window were identified as abnormal, the entire window was excluded from the HRV analysis. It has been indicated that, during an ultra-short recording period (< 5 minutes), the Root-Mean Square of Successive Difference (RMSSD) could be more reliable than the Standard Deviation of NN-interval (SDNN) (Baek et al. 2015; Munoz et al. 2015). We therefore calculated RMSSD as an indicator of the cardiac autonomic nervous system.

### Respiration

The respiration signals were band-pass filtered between 0.01 and 2 Hz before analysis. Outlier points were detected as data points exceeding 4 standard deviations. Trials with outlier ratio over 20% were excluded from analysis, otherwise, outlier points were linearly interpolated, based on the continuous hypothesis of physiological signals (Berger et al. 1986). Respiration peaks were identified using an adaptive prominence-based detection algorithm, as adopted in the vastly used toolbox NeuroKit2 (Makowski et al. 2021), where the minimum prominence of the peak was set at 25% of the 90% data range, calculated as the difference between the 95th and 5th percentiles, to capture inhalations while ignoring low-amplitude noise and motion artifacts. To ensure physiological plausibility, the peak-to-peak interval was constrained between 2.4 and 7.5 s, corresponding to a respiratory rate of 8–25 bpm. To ensure reliable respiration intervals, trials with fewer than 3 detected peaks were excluded. Also, the coefficient of variation (CoV = SD/mean) of the intervals, were calculated (Thamrin et al. 2022). Trials with high irregular respiration intervals (CoV > 15%) were excluded from the analysis.

### Galvanic skin response

The raw GSR signal was filtered with a low-pass filter of 1 Hz and converted from kΩ to micro-Siemens (μS). Out-of-range signals were identified when absolute values were lower than 0.5 μS or higher than 60 μS (Kleckner et al. 2018). Signal discontinuities were identified when rate of change between adjacent samples exceeded 10 μS. Trials containing out-of-range or discontinuity more than 30% of samples were excluded. For the remaining trials, samples identified as out-of-range or discontinuity were interpolated with *fillmissing.m*, where the points in the middle were linearly interpolated, and the points at each end were filled with nearest number. The interpolated data were then decomposed into tonic and phasic GSR signals with the cvxEDA algorithm (Greco et al. 2016). Tonic GSR signals from all trial collected on the same experimental day were normalized using z-score transformation. Trial with flat signals, defined as a SD < 0.001 μS, were excluded. The remaining trials were segmented into different window lengths to extract GSR indicators. Two GSR indicators were extracted from each trial, at each length: The tonic GSR level (skin conductance level, SCL) by averaging the normalized tonic signals; the phasic GSR activity (skin conductance component, SCC), indexed by the number of peaks. Peaks were defined by the MATLAB function *findpeaks.m* (minimum prominence = 0.1 SD and minimum peak distance = 1 second).

### Data quality check

After the preprocessing and artifact rejection, we check the trial retention in our dataset (result of trial rejection is illustrated in Supplementary Figure 1). During data inspection, we found one participant (Subject 9) showed atypical ECG morphology that precluded reliable RR-interval detection. Data from Subject 9 was therefore excluded from all subsequent analyses to ensure signal integrity. Furthermore, respiration rate was excluded from the main analysis due to the extremely high rejection rate above 50% across all windows. SpO2 was also excluded from analysis due to its primary role as safety monitor, and its lack of physiological variation (<1%).

### Window-wise regression analysis

To examine whether pre-trial physiological indicators were associated with subsequent creative performance, and to compare the relative contributions of the indicators, we constructed a linear mixed-effects model. The model included normalized RMSSD, HR, SCC, and SCL as fixed-effect predictors of *Regularized C*, while accounting for repeated observations across participants, experimental days, and tasks. The model was applied with biophysiological data covering various lengths of pre-stimulus periods (11–30 s) independently, which makes 20 independent analyses. The final regression model is listed below:

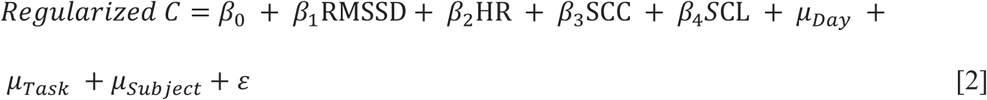

### Feature importance permutation test

Following the regression analysis, feature importance was evaluated separately for each window length using a permutation-based procedure. For each indicator, the correspondence between trials and that indicator was randomly shuffled within participant, thereby preserving the within-subject data structure while disrupting the association between the indicator and creativity performance. The same regression model was then fitted to the permuted dataset, and the reduction in model performance was calculated as the original model R^2^ minus the permuted model R^2^. This procedure was repeated 10,000 times for each indicator. Feature importance was quantified as the mean R^2^ reduction across permutations, and statistical significance was determined from the permutation distribution.

### Exploratory analysis of resting-state physiological dynamics

A previous analysis of EEG data collected in the same experiment identified a temporal property of the Creativity Potential Network (CPN; Chao, Street, and Liu 2025), a resting-state functional connectivity pattern associated with subsequent creativity performance. The CPN-related time series exhibited a mean dominant period of approximately 3.2 minutes, suggesting that the brain state associated with creativity potential may fluctuate over a relatively long timescale. We therefore conducted an exploratory follow-up analysis to examine the temporal correspondence between the physiological indicators and the previously reported CPN dynamics. Specifically, we estimated the dominant period of each physiological indicator and quantified its temporal distance from the group-level CPN reference period of 3.2 minutes.

To construct indicator-specific contribution time series, physiological features were extracted from each 5-min resting-state recording using a sliding window with the same duration as the optimal window identified in the primary regression analysis. The window was advanced in 1-s increments, and each feature was quantified using the same procedure as in the regression analysis. Feature values were standardized within participant and task, and the contribution of each physiological indicator was calculated as *Contribution*_k(t)_ = *β*_k_*z*_i_(*X*_k(t)_)), where *k(t)* indicates indicator *k* at window *t*, *β*_k_ indicates the corresponding coefficients at the optimal window, and *z*_i_(*X*_k(t)_) indicates the normalized feature value of indicator *k* within participant ***i.*** Missing values in the contribution time series were linearly interpolated when the proportion of missing windows did not exceed 20; recordings with more than 20% missing windows were excluded. Each contribution time series was then linearly detrend before entering next step analysis.

Dominant temporal periods estimated with power spectral density analyses, consistent with procedure used in the previous EEG study. PSD was calculated using the *periodogram.m* in MATLAB, and nonzero frequencies were converted to periods as (P = 1/f). To maintain methodological comparability, the maximum candidate period was determined by the duration of the 5-min recording. Periods shorter than 30 s were excluded to reduce the influence of short-timescale temporal dependence introduced by the highly overlapping sliding windows. The dominant period of each indicator was defined as the period longer than 30 s with the maximum spectral power. For each physiological indicator and resting-state recordings, temporal distance from CPN period was calculated as: *D*_k_ = 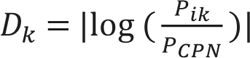 where *P_ik_* denotes the dominant period for indicator *k* in subject *i* for each resting recording. Distance 0 indicates a dominant period identical to CPN period, which equals 3.2 minutes.

## Results

### Normalization aligns GPT-rated creativity scores across tasks

Figure 3A shows the *C_AUT_* and *C_FIT_* scores across trials (questions) and participants. *C_AUT_* scores were approximately normally distributed, whereas *C_FIT_* scores were negatively skewed, with most observations concentrated at the higher end of the scale (Figure 3B). To account for these distributional differences, scores were normalized within each task and participant. The resulting *Regularized C* scores showed more comparable distributions across tasks (Figure 3C).

**Figure 3.**
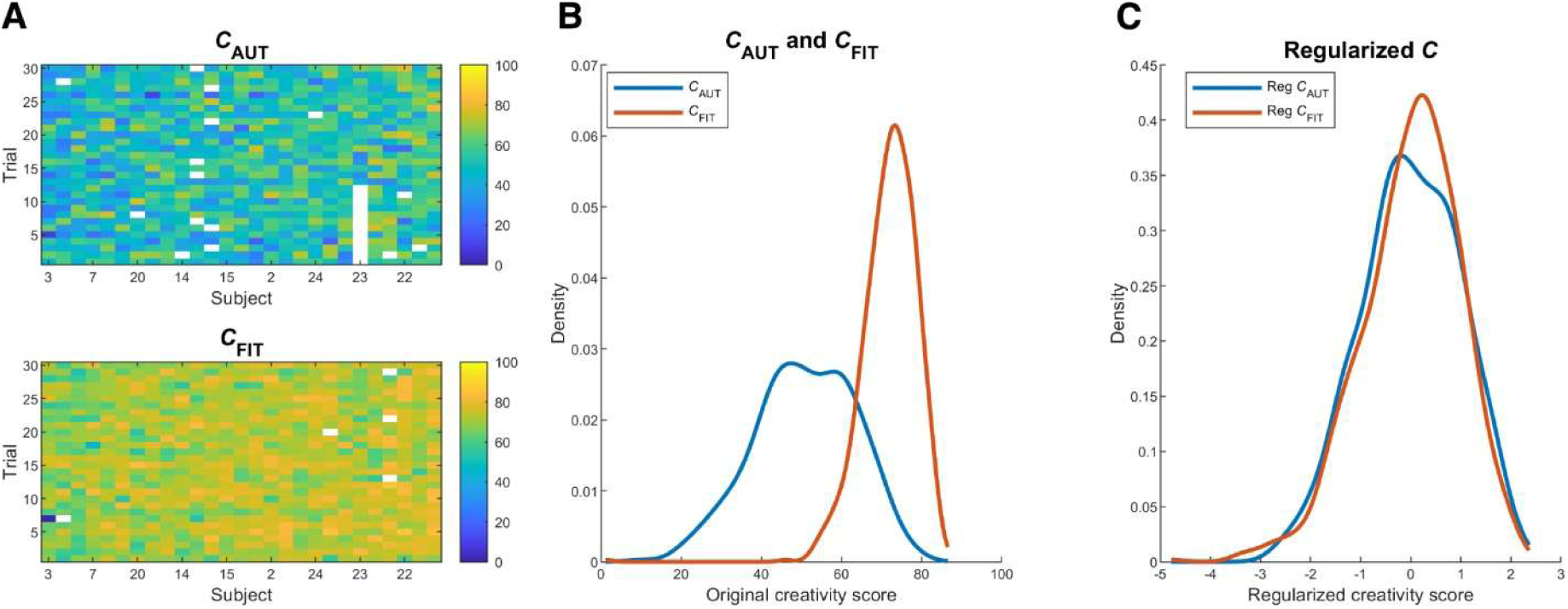
Creativity scores distributions aligned between tasks after normalization. (**A**) The heatmap demonstrated subject-wise and trial-wise *C_AUT_* (upper) and *C_FIT_* (lower), calculated from the geometric means of novelty and feasibility scores in AUT, and of novelty, feasibility, and goal attainment scores in FIT, with missing trials left blank. (**B**) The line plot demonstrates frequency distribution of *C_AUT_* (blue) and *C_FIT_* (orange). There is significant between-task difference that FIT showed right-skewed and sharp distribution over higher score range. To eliminate the task difference, all creativity was normalized within-subject and within-task by z-scores. (**C**) The *Regularized C* of the two tasks are aligned together, centered at 0.

### HR and SCC increased, but SCL decreased in longer observation windows

The number of trials included in the regression analysis at each window length is shown in Figure 4A. Most window-wise models included approximately 1,050 to 1,060 trials. The number of available trials was slightly lower for the shortest (11 s; 1043 trials) and longest (30 s; 1038 trials) windows, reflecting a greater proportion of trial for which valid physiological indicators could not be obtained due to limited data length, or more noise that cause trial exclusion.

**Figure 4.**
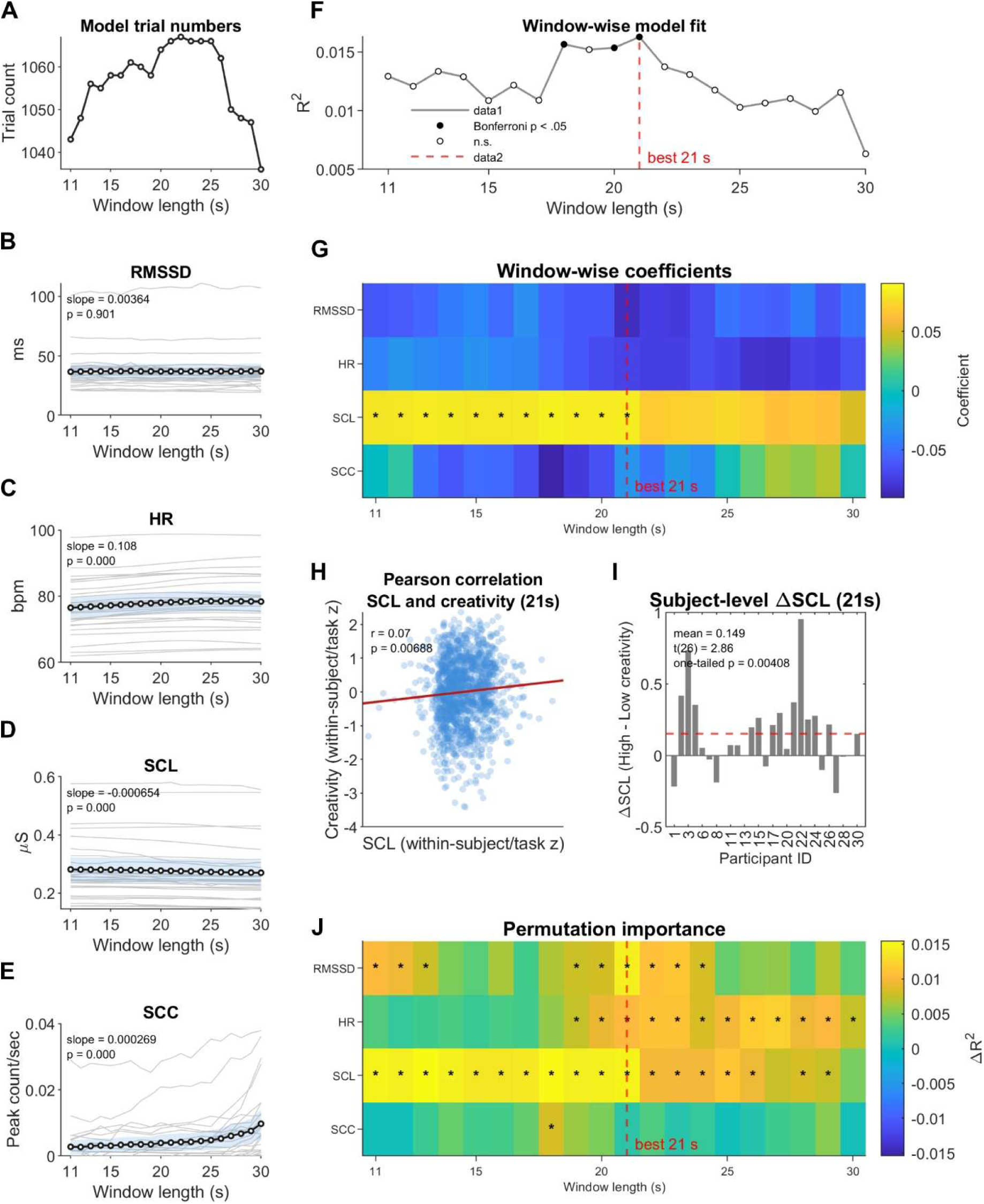
Physiological data descriptions and regression analysis results. (**A**) Survival trial numbers in window-wised regression models ranged between 1040-1070. From 11 s, model trial numbers increased with data length, reached higher level around 20-25 s, and dropped when data length exceeding 26 s. (**B-E**) Grand mean values and trends of RMSSD, HR, SCL and SCC in frequency (peak number/second). Black lines with circles indicate the grand mean values with blue shadings indicate 95% confidence intervals. Trends of each participant are plotted as bright gray lines. Slope of each indicator were examined by a permutation test of 5,000 iterations. The results indicated that RMSSD stayed stable across window length, however, HR (slope = 0.108, p <0.0001) and SCC rate (slope = 2.69e-04, p <0.0001) increased, while SCL (slope = −6.54e-04, p <0.0001) decreased in longer windows. (**F**) Line plot of window-wise model goodness of fit (standardized R-squared values) with model significance (p-values are Bonferroni-corrected) marked in black circles. The model showed a higher explanatory power with data length around 20 s and peaked at 21 s (R2 = 0.016, p = 0.0015), which is selected as the best observation window for later analysis. (**G**) The heatmap of window-wised standardized beta coefficients, with black asterisks marking FDR-corrected significance. SCL performed as a positive predictor to creativity scores, showing robust significance across window length shorter than 21 s. (**H**) Pearsons’ correlation between normalized SCL and *Regularized C* in the selected 21-second window showed a weak but robust positive correlation (r = 0.07, p = 0.0069). (**I**) Bar graph showing individual SCL differences (ΔSCL) between high- and low-creativity trials based on within-subject median split. The result showed positive ΔSCL in most participants and was positively greater than 0 (one-sample t-test: t26 = 2.86, p = 0.0041, Cohen’s d = 0.55). (**J**) Heatmap demonstrates feature importance in each data length, represented by mean R2 drops after shuffling each indicator 10,000 times in identical regression model. Permutation significance (p-values were FDR-corrected) was marked as black asterisks.

To assess the stability of each physiological indicator across window lengths, participant-level slopes were calculated across the 11–30 s windows and evaluated using permutation tests with 10,000 iterations. Figure 4B to 4E shows the individual participant trajectories and the group-average trend for each indicator. RMSSD remained stable across window lengths (Figure 4B, *slope* = 0.0036, *p* = 0.9), while HR increased slightly as window length increased (Figure 4C, *slope* = 0.108, *p* < 0.0001). Among the GSR indicators, SCL showed a small decrease with increasing window length (Figure 4D, *slope* = −0.0007, *p* < 0.0001), whereas SCC showed a small positive trend (Figure 4E, *slope* = 0.0003, *p* < 0.0001).

### SCL as the primary predictor of subsequent creativity performance

Window-wise fixed-effects regression models showed significant overall model fit at the 18-, 20-, and 21-s windows after Bonferroni correction (Figure 4F). Model fit was highest for the 21-s window (*R^2^* = 0.016), which was therefore selected as the optimal window for subsequent analyses. As shown in Figure 4G, SCL was a positive predictor of *Regularized C* from the 11- to 21-s windows after FDR correction. In contrast, the coefficients for RMSSD, HR, and SCC were not significant at any individual window length.

Together, the regression-coefficient and feature-importance analyses identify tonic SCL as the most consistent physiological indicator of creativity performance. At the optimal 21-s window, normalized SCL was positively correlated with *Regularized C* (*r* = 0.07, *p* = 0.0069; Figure 4H). To determine whether this association reflected a general within-subject tendency rather than being driven by a small number of extreme observations, each participant’s trials were divided into high- and low-creativity groups using a median split of *Regularized C*. Mean SCL was then calculated separately for high- and low-creativity trials, and a within-subject difference score was computed as high-creativity SCL minus low-creativity SCL. These difference scores were tested against zero using a one sample, one-tailed *t*-test. As shown in Figure 4I, most participants showed a positive difference, and mean SCL was significantly greater during high-creativity than low-creativity trials (*MΔ_SCL_* = 0.149, *t (26)* = 2.86, one-tailed *p* = 0.004). This result supports the interpretation that the positive SCL-creativity association was broadly observed across participants.

### Feature-importance analysis confirmed robust contribution from SCL

The results of permutation-base feature importance analysis are illustrated in Figure 4J, with asterisk marking FDR-corrected significance. The result suggested that permuting SCL substantially reduced model *R^2^* indicating that the explanatory performance of the regression model was impaired when the trial-wise SCL-creativity correspondence was disrupted.

RMSSD and HR also showed significant importance in selected windows, particularly around the optimal 21-s window. However, their regression coefficients were not statistically significant. Thus, although these cardiac indicators may contribute information to the fitted model, the current results do not provide evidence for a reliable independent linear association between RMSSD or HR and creativity.

### SCL exhibited a temporal scale closer to the CPN period compared to other indicators

Using the regression coefficients estimated from the optimal 21-s model, indicator-specific contribution time series were calculated from the 5-min resting-state recordings preceding each experimental session (Figure 5A). PSD analysis revealed dominant periods ranging from approximately 0.9 to 4.3 min across participants and indicators (Figure 5B). Among the four indicators, SCL showed the longest mean dominant period, with its group mean located near the previously reported CPN period of 3.2 min.

**Figure 5.**
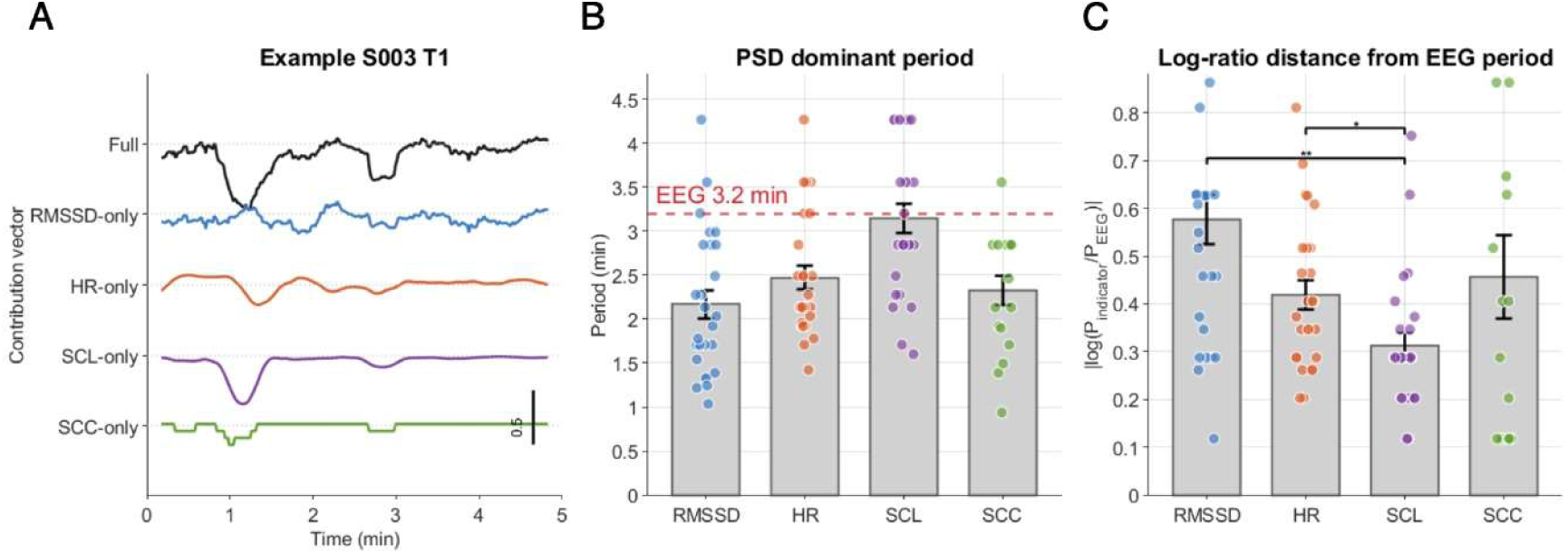
Exploitative analyses of indices dominant periods with the 5-minute resting data. (**A**) Example indicator-specific contribution time series from one resting recording. Physiological features were extracted using 21-s sliding windows advanced in 1-s steps, standardized within subject, and multiplied by the corresponding regression coefficient from the 21-s regression model. Each trace therefore represents the model-weighted partial contribution of one physiological indicator. (**B**)Dominant periods of the four indicator-specific contribution time series estimated by power spectral density (PSD) analysis. Gray bars and error bars indicate the grand mean and SEM, respectively, and colored dots indicate individual participant estimates. The dashed red line indicates the previously reported EEG dominant period of 3.2 min. Among the four indicators, the SCL-specific contribution showed a dominant period most comparable to the EEG period. (**C**) Distance between each physiological dominant period and the EEG dominant period. Distance was computed as the absolute log-ratio, |log(*P*_indicator_/*P*_EEG_)|, where P_EEG_ =3.2 min. Result-contingent contrast was made between distance of SCL and mean distance of RMSSD, HR, and SCC and showed significant result, suggesting that SCL has a dominant period closer to CPN. Pair-wise permutation demonstrates that dominant period of SCL was significantly more similar to CPN, compared RMSSD and HR, but not SCC.

To quantify temporal proximity to the CPN, the dominant period of each physiological indicator was converted to an absolute log-ratio distance from the 3.2-min EEG reference period (Figure 5C), with smaller values indicating greater proportional similarity. Visual speculation suggested that SCL, the only indicator significantly associated with creativity, had the smallest mean distance from the CPN period. We further conducted a permutation test (10,000 iterations) comparing its distance with the mean distance of other indicators.

Results suggested that period distance between SCL and CPN was significantly shorter than the average distance of RMSSD, HR, and SCC (one-tailed *p* = 0.001). Follow-up permutation tests compared SCL separately with each of the other indicators showed significantly smaller distance for SCL than for RMSSD (one-tailed *p* = 0.0005) and HR (one-tailed *p* = 0.008), but only marginally smaller than SCC (one-tailed *p* = 0.065). Therefore, SCL exhibited greater temporal similarity to the CPN period compared to other indicators on average, although this difference was not consistently significantly relative to every individual indicator.

## Discussion

The present study examined whether physiological signals recorded at rest contain information about momentary creative potential. We evaluated whether heart rate variability, indexed by the RMSSD, HR, SCL and SCC measured during a 30-s pre-trial resting period predicted subsequent performance on divergent-thinking (AUT) and convergent-thinking (FIT) tasks. Window-wise linear regression analyses indicated that these physiological measures had modest but statistically reliable information about subsequent creative performance, with model performance peaking at a window length of approximately 20 s. These findings provide preliminary evidence that spontaneous autonomic activity during a brief task-free period may reflect momentary physiological states associated with readiness for creative problem solving.

Among the physiological measures, tonic electrodermal activity, indexed by SCL, showed the most consistent association with creative performance. Higher pre-trial SCL was associated with better subsequent performance, particularly for analysis windows of up to 21 s.

Permutation-based feature-importance analysis further supported the contribution of SCL, as shuffling this indicator significantly reduced model explanatory performance. HR and RMSSD also contributed to overall model performance, although their coefficients were not significant, suggesting that these cardiac indicators may contain weak information relevant to model fit without showing stable associations with creativity. Thus, the most consistent evidence across analyses was obtained for SCL.

Electrodermal activity (EDA), also referred to as galvanic skin response (GSR), reflects changes in the electrical conductance of the skin resulting from sweat-gland activity. Because eccrine sweat glands are primarily controlled by the sympathetic nervous system, EDA is commonly used as an index of sympathetic arousal. Our findings therefore suggest a positive association between pre-trial tonic sympathetic arousal and subsequent creative performance. This result is consistent with previous studies linking task-related arousal to creativity (Loudon and Deininger 2016; Shen et al. 2018) while extending this association to physiological activity measured before task engagement. Because SCL was assessed during a task-free pre-trial period, the observed relationship is less likely to reflect an immediate response elicited by the creative task itself. Instead, it may reflect spontaneous fluctuations in autonomic state that contribute to moment-to-moment variation in readiness for creative thinking.

Building on EEG findings from the same experiment (Chao et al., 2025), we conducted an exploratory analysis of the intrinsic temporal scales of the physiological indicators by estimating their dominant periods during a 5-min resting-state recording. Using the optimal 21-s observation window identified in the regression analysis, we found that the physiological indicators exhibited fluctuations on the scale of minutes. Notably, SCL, the only physiological indicator that significantly predicted subsequent creative performance, showed a dominant period closer to that of the creativity potential network (CPN), a connectivity-based EEG marker of creativity, than did the non-significant physiological indicators. This raises the possibility that they track complementary peripheral and cortical manifestations of a common, slowly evolving psychophysiological state associated with readiness for creative thinking.

One candidate mechanism underlying this slowly evolving state is the locus coeruleus– noradrenergic (LC-NA) system, which broadly regulates cortical arousal, neural gain, behavioral engagement, and cognitive flexibility (Arnsten and Goldman-Rakic 1984; Xing, Li, and Gao 2016; Berridge and Spencer 2016; Aston-Jones and Cohen 2005). According to Adaptive Gain Theory, changes in tonic LC activity modulate the balance between an exploitative mode, characterized by focused task engagement, and an exploratory mode, characterized by greater responsiveness to alternative or weakly represented information (Aston-Jones and Cohen 2005). Consistent with this account, animal studies have shown that elevated LC activity, whether spontaneous or optogenetically induced, can facilitate rule switching, a behavioral index of cognitive flexibility (McBurney-Lin et al. 2022). In humans, studies using pupil diameter as a noninvasive proxy of LC-related arousal have reported associations between pre-trial tonic activity and originality in both standardized divergent-thinking tasks and more naturalistic creative tasks, although the evidence is not fully consistent across studies (de Rooij, Vromans, and Dekker 2018; also see Jos et al. 2025). On this basis, the slow fluctuations in LC-NA activity may influence both peripheral sympathetic arousal and large-scale cortical dynamics, which may be reflected in SCL and CPN, respectively. However, this account remains provisional because neither measure directly indexes LC activity nor similarity in temporal scale alone cannot establish a common physiological origin.

Although SCL may reflect a physiological state associated with creativity, it is also sensitive to broader variations in arousal and task engagement. The observed association between SCL and creative performance could therefore reflect greater engagement, faster ideation, or lower fatigue rather than a creativity-related state per se. To evaluate these alternative explanations, we added normalized response time, z-scored within participant and task, to the regression model as an index of response speed and task engagement. We also included trial order within each session to account for systematic changes over time, including fatigue, habituation, and practice effects. Three additional models were fitted by adding trial order, normalized response time, or both variables to the original model.

We then examined whether inclusion of these covariates altered the optimal window, reduced the magnitude or significance of the SCL coefficient, or increased collinearity involving SCL. As shown in Supplementary Figure 4, the optimal 21-s window remained unchanged across all models. The magnitude of the SCL coefficient increased rather than decreased after adjustment, and its range of statistical significance was preserved. Although the variance inflation factor for SCL increased slightly, it remained close to 1, indicating negligible collinearity. These findings indicate that the association between pre-trial SCL and subsequent creative performance was not readily explained by response speed or by systematic trial-related changes such as practice, habituation, or fatigue. The persistence of the SCL effect after covariate adjustment therefore supports the robustness of the primary result.

Several limitations should be considered. First, SCL is a nonspecific index of sympathetic arousal that is influenced by attention, stress, emotion, and task engagement. Although the association between SCL and creativity remained after adjustment for response time and trial order, these analyses cannot distinguish creativity-related readiness from other unmeasured affective or motivational states. Future studies should therefore include subjective ratings of arousal, mood, and engagement, together with complementary physiological measures such as pupil diameter, to clarify the processes underlying the SCL effect.

Second, the observed positive association was identified during a low-demand pre-trial resting period. It remains unclear whether the same relationship would generalize to conditions of moderate or high arousal. Previous work suggests that moderate physical activity may facilitate creativity, whereas the effects of more intense arousal are less consistent (Zhao et al. 2022; Rominger et al. 2022; also see Rominger et al. 2026 for review). Experimental manipulation of arousal through exercise or environmental conditions may help determine the range over which SCL remains positively associated with creative performance.

Third, the dominant-period analysis should be regarded as exploratory. Because the indicator-specific time series were calculated using a 21-s sliding window, the period search was restricted to periods longer than 30 s to reduce the possibility that the estimated temporal scales merely reflected the analysis-window duration. Nevertheless, a 5-min recording remains too short to characterize a reliable fluctuation with a period of approximately 3 min, as it contains fewer than two complete cycles. The estimated dominant periods should therefore be interpreted as descriptions of characteristic temporal scale rather than evidence of stable physiological oscillations. In addition, the comparison between the physiological indicators and CPN was conducted at the group level and does not establish within-participant coupling or synchronization. Longer simultaneous EEG and physiological recordings, followed by participant- and trial-level analyses, will be required to determine whether the peripheral and cortical signals exhibit coordinated temporal dynamics.

Finally, although the association between SCL and subsequent creative performance was statistically reliable across several robustness analyses, the model explained only a small proportion of behavioral variance, with an R^2^ of approximately 1.6%. Physiological state therefore appears to capture only one component of the substantial moment-to-moment variability in creative performance. The present findings should consequently be interpreted as evidence that pre-trial physiology contains modest information about subsequent creativity, rather than as support for accurate individual-level behavioral prediction.

In conclusion, the present study provides evidence that physiological activity during a brief pre-trial resting period contains modest information about subsequent creative performance across divergent- and convergent-thinking tasks. Among the examined autonomic indicators, SCL showed the most consistent association with creativity, suggesting that momentary tonic sympathetic arousal may inform the physiological state preceding successful creative thinking. Although the limited explained variance precludes accurate individual-level prediction, these findings establish a basis for further investigating whether wearable physiological signals can contribute to characterizing creativity-related states.

## Author contributions

Z.C.C. conceptualized the study. T.L.L. and M.S. implemented the experiment and collected the data. T.L.L. performed data analysis and wrote the paper. Z.C.C. and M.S. edited the paper. All authors contributed to and approved the final paper.

## Acknowledgments

This study was supported by the World Premier International Research Center Initiative (WPI), MEXT, Japan (to Z.C.C.), and the IRCN–Daikin SCP (to Z.C.C.).

## Supplementary Figures

**Supplementary Figure 1.**
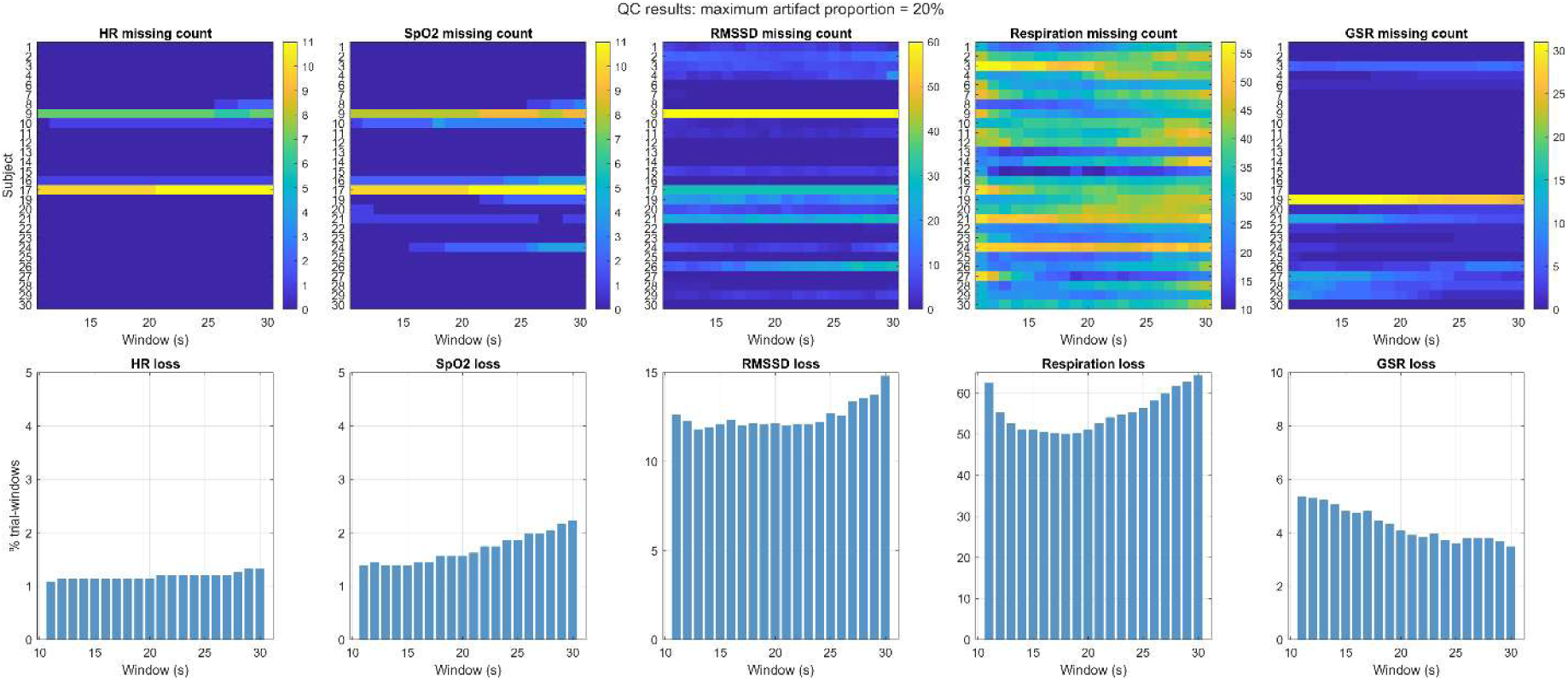
Evaluation of data quality and trial loss across window lengths. (Upper row) Heat maps indicating the frequency of missing trials per participant (listed along y-axis) across window length (x-axis). Subject 9 exhibited high data loss in the RMSSD due to atypical ECG morphology that precluded reliable RR-interval detection. All data from this participant was excluded from the following analysis. (Lower row) Bar graphs illustrate ratio of trial loss for each indicator in each window length. While data loss remained stable and low for HR, SpO2, HRV, and GSR, respiration data exceeded a 50% loss across all windows. Due to this insufficient signal reliability, respiration rate was omitted from the subsequent statistical analysis. Additionally, while data of SpO2 had very little data loss, it was also excluded from the main analysis, due to its role as a safety monitor, rather than an experimental indicator.

**Supplementary Figure 2.**
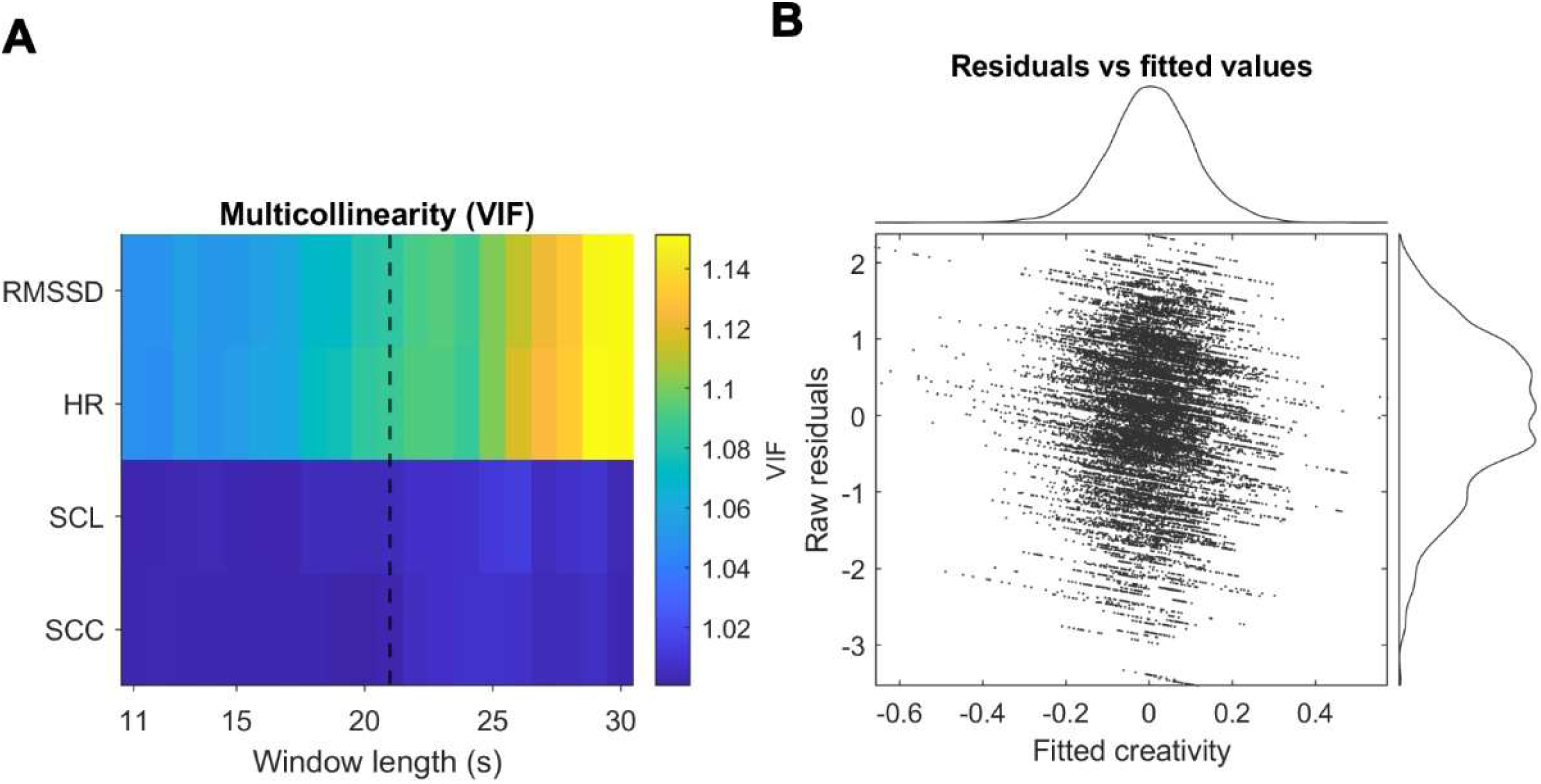
Diagnostic data of the regression models. (**A**) The multi-colinearity heat map of all predictors across window lengths suggested that colinearity values across all window length were close to 1. (**B**) Distribution of model residuals showed that the residuals were normally distributed and centered around zero, suggesting no violation on the regression hypothesis.

**Supplementary Figure 3.**
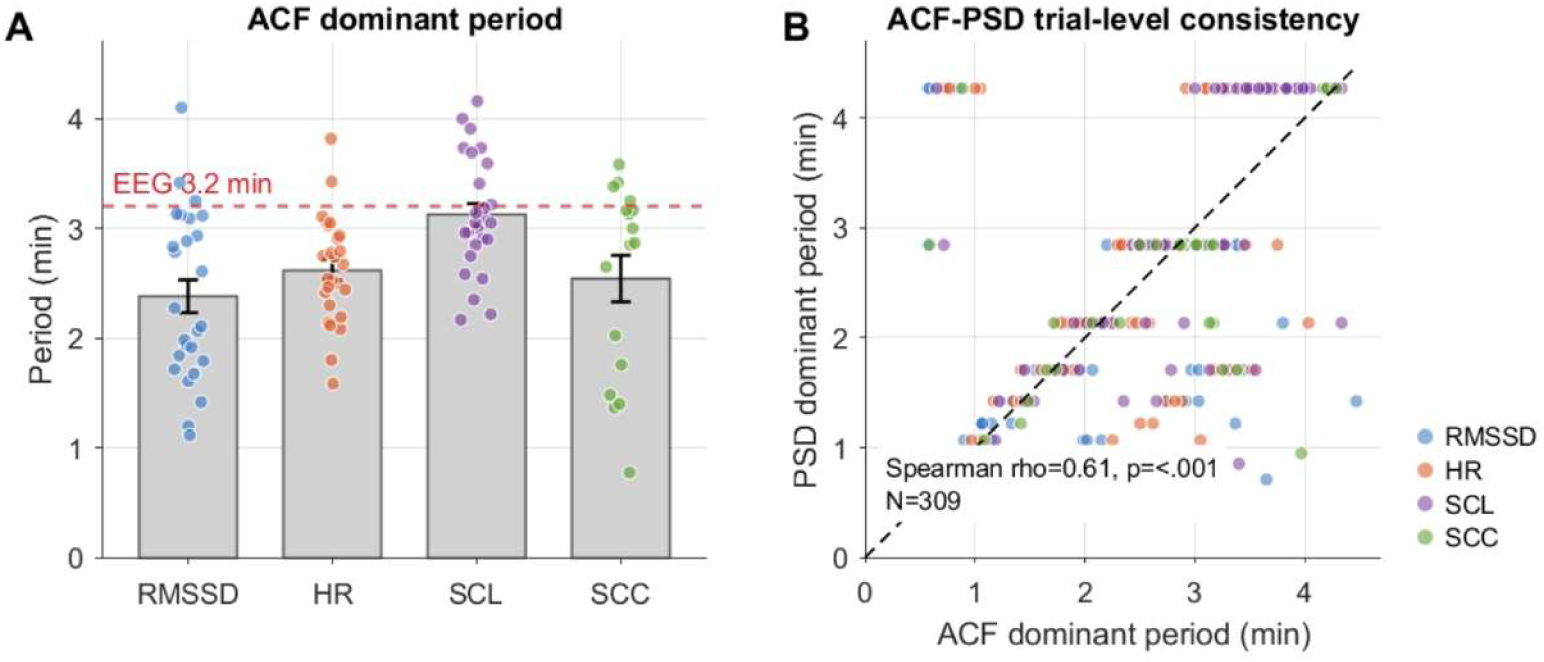
Convergence between ACF- and PSD-derived dominant period estimates. (**A**) Dominant periods of the four indicator-specific contribution time series estimated using autocorrelation function (ACF) analysis. Gray bars and error bars indicate the grand mean and SEM, respectively, and colored dots indicate individual recording estimates. The dashed red line indicates the previously reported EEG dominant period of 3.2 min. The ACF-derived pattern was broadly consistent with the PSD-derived dominant periods reported in the main analysis, with the SCL-specific contribution showing the dominant period closest to the EEG period. (**B**) Trial-level correspondence between dominant periods estimated by ACF and PSD. Each dot represents one estimated dominant period of one indicator from one trial. Spearman correlation showed a positive association between ACF- and PSD-derived dominant periods (Spearman *ρ* = 0.61, *p* < 0.001), supporting the convergence of the two temporal-scale estimation methods. This result suggests that the PSD-derived dominant periods used in the main analysis were not specific to a single estimation procedure.

**Supplementary Figure 4.**
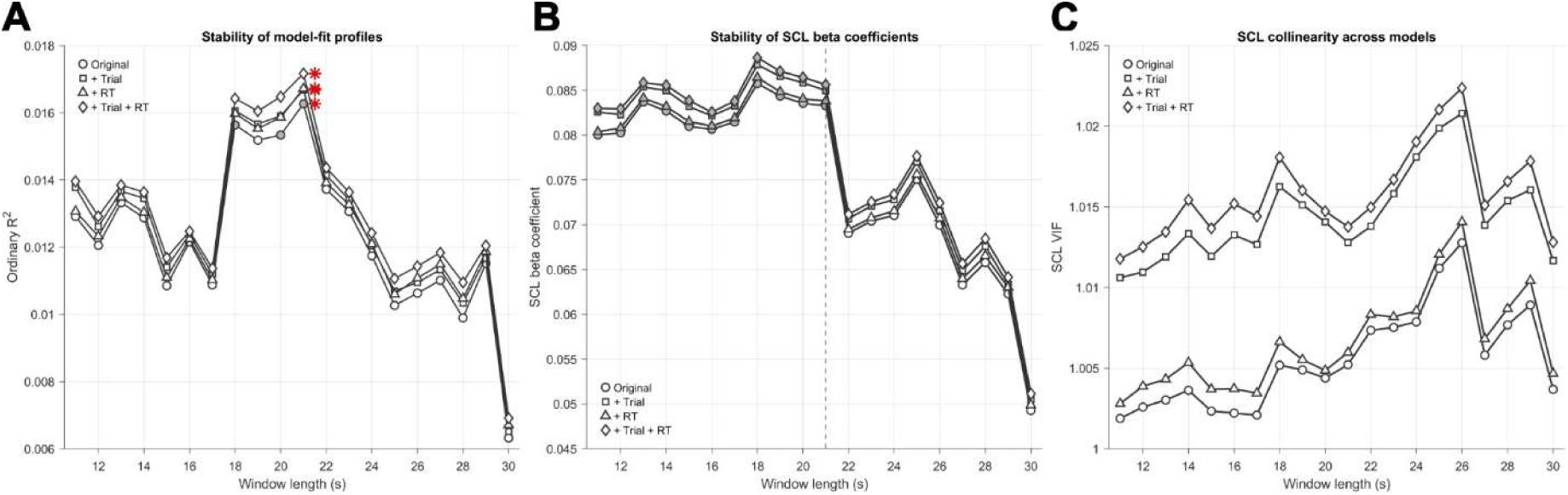
Examination of model sensitivity to reaction time and trial order. (**A**). Line plots of window-wise *R^2^* values across window lengths in original model (circle marker) and models including trial order (Trial, square marker), response time (RT, triangle marker), or both RT and Trial (diamond marker). Significant models after the Bonferroni correction were informed with solid markers, the best windows with peak *R^2^* were marked as red asterisks next to the marker. The *R^2^* curves were highly overlapped between original and adjusted models. Best windows also agreed between models, remained at 21s. (**B**) Window-wise SCL beta coefficients showed that including RT and Trial did not decrease (but slightly increased) the magnitude or significance of the SCL beta coefficients, suggesting that the SCL-creativity relation was robust. (**C**) Variance inflation factors for SCL across models and window lengths, showing negligible collinearity after covariate adjustment.

